# Microglial expression of the Wnt signalling modulator *DKK2* differs between human Alzheimer’s disease brains and mouse neurodegeneration models

**DOI:** 10.1101/2022.05.05.490649

**Authors:** Nozie D. Aghaizu, Sarah Jolly, Satinder K. Samra, Bernadett Kalmar, Katleen Craessaerts, Linda Greensmith, Patricia C. Salinas, Bart De Strooper, Paul J. Whiting

## Abstract

Wnt signalling is crucial for synapse and cognitive function. Indeed, deficient Wnt signalling is causally related to increased expression of *DKK1,* an endogenous negative Wnt regulator, and synapse loss, both of which likely contribute to cognitive decline in Alzheimer’s disease (AD). Increasingly, AD research efforts have probed the neuroinflammatory role of microglia, the resident immune cells of the central nervous system (CNS), which have furthermore been shown to be modulated by Wnt signalling.

The *DKK1* homologue *DKK2* has been previously identified as an activated response and/or disease-associated microglia (DAM/ARM) gene in a mouse model of AD. Here we performed a detailed analysis of *DKK2* in mouse models of neurodegeneration, and in human AD brain. In *APP/PS1* and *APP^NL-G-F^* AD mouse model brains as well as in *SOD1^G93A^* ALS mouse model spinal cords, but not in control littermates, we demonstrated significant microgliosis and microglial *Dkk2* mRNA upregulation in a disease-stage dependent manner. In the AD models, these DAM/ARM *Dkk2^+^* microglia preferentially accumulated close to βAmyloid plaques. Furthermore, recombinant DKK2 treatment of rat hippocampal primary neurons blocked WNT7a-induced dendritic spine and synapse formation, indicative of an anti-synaptic effect similar to that of DKK1. In stark contrast, no such microglial *DKK2* upregulation was detected in the post-mortem human frontal cortex from individuals diagnosed with AD or pathological ageing.

In summary, the difference in microglial expression of the DAM/ARM gene *DKK2* between mouse models and human AD brain highlights the increasingly recognised limitations of using mouse models to recapitulate facets of human neurodegenerative disease.

**Significance statement:** The endogenous negative Wnt regulator *Dkk2* is significantly upregulated at the mRNA level in microglia of AD mouse models, implying that microglia derived Dkk2 protein may detrimentally contribute to a reduced Wnt signalling tone in the AD brain, a known pathophysiological manifestation. Indeed, recombinant DKK2 prevented Wnt-dependent synapse formation in cultured neurons. However, *DKK2* upregulation was not recapitulated in post-mortem human AD brains.

The success of neurodegeneration animal models has relied on pathophysiology that for the most part correctly modelled human disease. Increasingly however, limitations to the validity of mouse models to recapitulate human neurodegenerative disease have become apparent, as evidenced by the present study by the difference in microglial *DKK2* expression between AD mouse models and human AD brain.

## Introduction

Microglia, the resident immune cells of the central nervous system (CNS), contribute both beneficially and detrimentally to Alzheimer’s disease (AD) in a context-dependent manner, thus rendering their response to AD heterogeneous in nature. So too is their transcriptomic and proteomic phenotype, leading to the identification of spatio-temporally distinct microglial subpopulations (reviewed in Masuda et al., 2020). Disease associated (DAM) – or activated response microglia (ARM; henceforth: DAM/ARM) – represent a subpopulation associated with the neurodegenerative brain (Keren-Shaul et al., 2017; Sala Frigerio et al., 2019). Transitioning from resting to DAM/ARM-state requires TREM2 (triggering receptor expressed on myeloid cells-2) (Keren-Shaul et al., 2017). A bona fide receptor for βAmyloid, TREM2 ligation activates microglia and orchestrates a gene regulatory response that increases inflammatory signalling, phagocytosis, and proliferation, a response thought to restrict development of AD (reviewed in Gratuze et al., 2018).

TREM2 regulates microglial proliferation and survival by activating, among others, the canonical Wnt/β-catenin pathway (Zheng et al., 2017; reviewed in Aghaizu et al., 2020). Indeed, several genes upregulated by TREM2 in response to AD pathology are related to proliferation and Wnt signalling (Meilandt et al., 2020). The canonical Wnt signalling modulatory gene *Dkk2* (Mao and Niehrs, 2003) was upregulated downstream of *Trem2* in DAM/ARM cells in *APP/PS1*, *PS2APP, 5xFAD, and APP^NL-G-F^* AD mouse models in separate studies, making it a putative DAM/ARM marker gene (Friedman et al., 2018; Sala Frigerio et al., 2019; Meilandt et al., 2020). The secreted protein DKK2 belongs to the Dickkopf family of Wnt modulators (Niehrs, 2006). Its homologue DKK1 antagonises Wnt signalling through Frizzled Wnt receptors by sequestering the co-receptor LRP5/6 (Bafico et al., 2001; Mao et al., 2001). The reduced Wnt signalling tone evident in AD is at least partially due to Aβ fibril-induced upregulation of *DKK1/Dkk1* in human AD and AD mouse models (Caricasole et al., 2004; Rosi et al., 2010; Killick et al., 2014; Sellers et al., 2018; Jackson et al., 2019). This was synaptotoxic in *in vitro* and *in vivo* models (Purro et al., 2012; Galli et al., 2014; Marzo et al., 2016; Elliott et al., 2018; Sellers et al., 2018), and potentially also in human AD (Jackson et al., 2019).

Much less is known about the role of microglial *DKK2* in the CNS, not to mention in AD. In cell lines, DKK2 can both antagonise and agonise Wnt-LRP6 signalling depending respectively on the presence or absence of the second co-receptor Kremen2 (Mao and Niehrs, 2003). During neural crest specification, DKK2 agonises Wnt signalling (Devotta et al., 2018). Conversely, in cancer studies DKK2 generally inhibits Wnt signalling (Kuphal et al., 2006; Sato et al., 2007; Maehata et al., 2008; Hirata et al., 2009; Zhu et al., 2012; Mu et al., 2017). Furthermore, cancer cell-secreted DKK2 suppresses immune cell activation via an unconventional Wnt- unrelated pathway (Xiao et al., 2018). In the aforementioned single-cell and bulk cell gene expression studies on neurodegeneration mouse models, *Dkk2* was upregulated in microglia, but no information on the spatial relationship between *Dkk2^+^* microglia and neurodegenerative pathology or the biological role of this upregulation was provided (Friedman et al., 2018; Sala Frigerio et al., 2019; Meilandt et al., 2020). To address this gap in our knowledge, we performed a histological assessment of microglial *Dkk2/DKK2* upregulation in several mouse models and in human AD, and furthermore investigated the effect of recombinant DKK2 on cultured primary neurons.

Here, we report significant microgliosis and microglial *Dkk2* mRNA upregulation in a disease-stage dependent manner in *APP/PS1,* and *APP^NL-G-F^* AD mouse model brains. Clustering of *Dkk2*^+^ microglia around amyloid plaques was often more pronounced than that of *Dkk2*^-^ microglia. In cultured rat neurons, recombinant DKK2 blocked Wnt dependent synapse formation. Crucially however, microglial *DKK2* upregulation was not detected in post-mortem human brain from individuals diagnosed with AD or pathological ageing. This non-universality of what was a putative DAM/ARM marker gene highlights the increasingly recognised limitations of using animal models to recapitulate facets of human neurodegenerative disease.

## Materials and Methods

### Mice

Mouse CNS tissue was obtained from the following sources: Brain tissue from male and female *B6.Cg-Tg*(*APPswe, PSEN1dE9*) mice (Jankowsky et al., 2004; RRID:MMRRC_034829-JAX; abbreviated: *APP/PS1*) and age-matched wild-type *C57BL/6J* control mice at 3, 8, and 12 months was purchased from WuXi AppTec. Brain tissue from male homozygous *B6.129S5-Apptm3.1Tcs* mice (Saito et al., 2014; RRID:IMSR_RBRC0634; backcrossed for at least 2 generations with *C57BL/6J* mice; here referred to as *APP^NL-G-F^*) and age-matched wild-type *C57BL/6J* control mice at 7 and 24 months was kindly provided by the De Strooper lab. Spinal cord tissue from female mice expressing mutant human *SOD1^G93A^*(*B6SJL*-*Tg[SOD1^G93A^]1Gur/J*; Gurney et al., 1994; RRID:IMSR_JAX:002726; abbreviated: *SOD1^G93A^*) and age-matched female control mice expressing wild-type human *SOD1* (*B6SJLTg[SOD1]2Gur/J*; Gurney et al., 1994; RRID:IMSR_JAX:002297; abbreviated: *SOD1^WT^*) at 50, 100, and 120 days was kindly provided by the Greensmith lab. Colonies were maintained by breeding male heterozygous carriers with female (*C57BL/6 × SJL*) F1 hybrids. Mice were genotyped for the human *SOD1* transgene from ear or tail genomic DNA.

In every case, mice were housed according to the appropriate institution’s ethical requirements, and in compliance to the country’s laws for animal research. Typically, mice were housed in standard individually ventilated cages with ≤ 3 mice per cage at 21±1 °C with relative humidity 55±10 % and maintained on a 12-hour light/dark cycle with access to food (standard pellets), water, and nesting material provided *ad libitum* via an overhead rack. At the onset of pathology, affected animals were provided with food pellets soaked in water at ground level to ensure sufficient nourishment and hydration. Cages were checked daily to ensure animal welfare. Body weight was assessed regularly to ensure no weight loss. For animals housed at WuXi AppTec, studies were reviewed and approved by Institutional Animal Care and Use Committee (IACUC) of WuXi AppTec (Suzhou) Co., Ltd. For animals housed at VIB/KU Leuven, studies were approved by the KU Leuven Ethical Committee and in accordance with European Directive 2010/63/EU. For animals housed at UCL, studies were carried out following the guidelines of the UCL Institute of Neurology Genetic Manipulation and Ethic Committees and in accordance with the European Community Council Directive of November 24, 1986 (86/609/EEC). Animal experiments were undertaken under licence from the UK Home Office in accordance with the Animals (Scientific Procedures) Act 1986 (Amended Regulations 2012) and were approved by the Ethical Review Panel of the Institute of Neurology.

For tissue collection, animals were injected with terminal anaesthesia (pentobarbital sodium, Euthatal) and were transcardially perfused with PBS by trained personnel.

### Rats

Animal experiments were undertaken under licence from the UK Home Office in accordance with the Animals (Scientific Procedures) Act 1986 (Amended Regulations 2012) and in compliance with the ethical standards at University College London (UCL). Timed matings were set up for Sprague-Dawley rats (RRID:MGI:5651135) for subsequent harvesting of embryos at embryonic day (E)18. Pregnant rat dams were sacrificed using Isoflurane and cervical dislocation.

### Human post-mortem tissue

Anonymised human samples from control, pathological ageing, and AD subjects were obtained from the Queen’s Square Brain Bank for Neurological Disorders (QSBB) and NeuroResource, UCL Institute of Neurology, University College London. All samples were obtained with informed consent in accordance with the Human Tissue Act 2004 and under the UCL Institute of Neurology HTA material transfer agreement UCLMTA1/17 approved by the NHS Research Ethics Committee. Post-mortem frontal cortex biopsy tissue was harvested, snap-frozen, and stored at -80 °C until further tissue processing. All experiments were performed in accordance with relevant guidelines and regulations. Sample information including demographic data, disease classifications and post-mortem intervals is shown in Supplementary Table S2.

### Tissue processing

Mouse brain tissue, freshly harvested upon transcardial perfusion with PBS, was post-fixed by immersion in 4 % paraformaldehyde (PFA) in PBS overnight at 4 °C followed by overnight immersion in and equilibration to 20 % sucrose in PBS at 4 °C. Mouse brains were split into 3 segments by applying 2 equidistant coronal slice cuts along the rostro-caudal axis, resulting in an olfactory bulb containing rostral-most segment, a hippocampus containing middle segment and the caudal-most cerebellar segment. After embedding in OCT (CellPath) and freezing in 2-methylbutane (Sigma) pre-chilled in liquid Nitrogen, the middle segment was coronally cryosectioned at 15 µm thickness on a Leica CM1860UV cryostat (Leica) and sections containing clearly defined hippocampus were transferred onto Superfrost Plus Gold microscopy slides (ThermoScientific).

Mouse spinal cord tissue was processed identically but split only into 2 segments by applying a transverse slice cut rostral to the lumbar enlargement, resulting in a rostral cervical/thoracic segment and a caudal lumbar segment. The cryo-embedded lumbar segment was transversally cryosectioned at 15 µm thickness and sections containing clearly defined L5 lumbar spinal cord were transferred onto Superfrost Plus Gold microscopy slides (ThermoScientific).

Human frontal cortex brain tissue was cryosectioned at 15 µm thickness, sections were transferred onto Superfrost Plus Gold microscopy slides (ThermoScientific) and dried for 10 min at 40 °C.

Human and mouse sections were stored at -80 °C until staining.

### mRNA fluorescence *in situ* hybridisation (FISH)

mRNA fluorescence *in situ* hybridisation for mouse *Dkk2* and human *DKK2*, *TREM2*, and *P2RY12* mRNA was performed on mouse brain / spinal cord and human frontal cortex cryosections respectively, by using the Multiplex Fluorescent V2 Assay Kit (ACD Bio).

Briefly, mouse cryosections were thawed and dried at 40 °C for 4 min prior to post-fixation with 4 % PFA in PBS at room temperature (RT) for 10 min. OCT residue was washed off by applying 1 x PBS for 5 min at RT. Sections were treated with RNAScope H2O2 for 4 min at RT and subsequently washed 2 x 3 min with UltraPure Distilled Water (Invitrogen) at RT. Microscopy slides containing cryosections were submerged for 4 min in boiling 1 x RNAScope target retrieval solution followed by immediate submersion in UltraPure Distilled Water. Cryosections were dehydrated in 100 % ethanol at RT for 2 min and allowed to air dry at RT for 5 min. Cryosections were subsequently treated with RNAScope Protease IV at RT for 15 min and washed 2 x 3 min at RT with 1 x PBS. RNAScope probes were allowed to hybridise to cryosections for 2 hrs at 40 °C (*Mm*-*Dkk2*-C1, 404841; *Mm-Ppib*-C1 (positive control probe), 313911; *E. coli*-*Dapb*-C1 (negative control probe), 310043). Probes were detected with TSA- Cy3 (Perkin Elmer, FP1170) using the RNAScope branched DNA amplification principle as per the manufacturer’s instructions. Subsequently, cryosections were further immunohistochemically processed (see below).

Human cryosections were processed similarly as previously described (Jolly et al., 2019). Briefly, cryosections were thawed and dried at 40 °C for 4 min prior to post-fixation with chilled 4 % PFA in PBS at 4 °C for 30 min followed by 2 x 2 min washes with 1 x PBS at RT. Cryosections were then dehydrated in an ethanol dilution series (50 %, 70 %, 2 x 100 %) at RT for 5 min each and subsequently allowed to air dry at RT for 5 min. Sections were treated with RNAScope H2O2 for 10 min at RT and subsequently washed 2 x 2 min with 1 x PBS. Microscopy slides containing cryosections were submerged for 10 min in boiling 1 x RNAScope target retrieval solution followed by 2 x 2 min washes with 1 x PBS. Cryosections were subsequently treated with RNAScope Protease IV at RT for 20 min and washed 2 x 3 min at RT with 1 x PBS. RNAScope probes were allowed to hybridise to cryosections for 2 hrs at 40 °C (*Hs*-*TREM2*-C1, 420491; *Hs*-*DKK2*-C2, 531131-C2; *Hs*-*P2RY12*-C3; 450391-C3; *Hs*-

*PPIB*-C1 (positive control probe), 313901; *E. coli*-*Dapb*-C1 (negative control probe), 310043). C2 and C3 probes were diluted in C1 probe solution at a 1:50 ratio. Probes were detected with TSA-Cy3 (Perkin Elmer, FP1170), Opal 620 (Akoya, FP1495001KT), and TSA-Cy5 (Perkin Elmer, REF FP1168) using the RNAScope branched DNA amplification principle as per the manufacturer’s instructions. Subsequently, cryosections were further immunohistochemically processed (see below).

### Primary hippocampal neuron cultures

Primary rat hippocampal neuron cultures were prepared from embryonic day 18 (E18) Sprague-Dawley rat embryos. One day prior to neuron isolation, 8 well chamber slide dishes (Miltenyi Biotec) were coated over night with 1 mg/ml poly-L-lysine in borate buffer (boric acid, 3.1 g/l; borax 4.8 g/l; pH 8.5). On the day of the neuron isolation, dishes were washed 3 x 20 min with UltraPure Distilled Water, filled with plating medium (Neurobasal (ThermoFisher) supplemented with 1x B27 (ThermoFisher), 1x GlutaMAX (ThermoFisher), 1x Penicillin-Streptomycin (ThermoFisher), 25 µM L-glutamate (Sigma)), and pre-equilibrated at 5 % CO2, 37 °C. Hippocampi were dissected from brain tissue using sterilized tools (Dumont #5 fine tip tweezers, Dumont #7 curved forceps, Student Vannas Scissors 9cm long/straight; Fisherbrand) and collected in ice cold HBSS (Invitrogen). Following three washes with fresh ice cold HBSS, hippocampi were enzymatically dissociated by incubation in accutase (ThermoFisher) at 37 °C for 10 min, providing manual agitation every 2-3 min. Hippocampi were then washed three times with pre-warmed (37 °C) HBSS, followed by mechanical dissociation into a single cell suspension by trituration in HBSS using a 1 mL pipette. Live cell density was determined using the Countess 3 automated cell counter (ThermoFisher) and cells were plated onto 8 well chamber slides at a density of 43,000 cells/cm^2^ and cultured in an incubator at 37 °C / 5 % CO2. Half medium changes were performed twice per week with maintenance medium: Neurobasal, supplemented with 1x B27, 1x GlutaMAX, 1x Penicillin-Streptomycin.

Neuronal transfection with the DNA construct *pHR hsyn:EGFP* (Keaveney et al., 2018; kind gift from Xue Han (Addgene plasmid # 114215; http://n2t.net/addgene:114215; RRID:Addgene_114215)) was performed at 7 days *in vitro* (DIV) using the Neuromag magnetofection method (OzBiosciences). Briefly, for every 40,000 cells plated per well of an 8 well chamber slide dish, 0.5 µg DNA was mixed and complexed with 1 µl Neuromag transfection reagent in 100 µl of OptiMem (all reagents at room temperature). Following 20 min incubation at room temperature, the transfection mix was added dropwise to neuronal cultures and the culture dish was placed on a magnetic plate (OzBiosciences) pre-equilibrated to 37 °C inside an incubator for the magnetofection step. After 20 min of magnetofection in the incubator, cell culture dish was removed from the magnetic plate and normal cell culture resumed.

Recombinant protein treatment was performed at 21 DIV for 24 hours: human DKK2 (Bio-Techne, 6628-DK-010/CF, 100 ng/ml), human DKK1 (Bio-Techne, 5439-DK-010/CF, 100 ng/ml), human WNT7a (Bio-Techne, 3008-WN-010/CF, 200 ng/ml); 100 ng/ml bovine serum albumin (BSA) in 1x PBS heat inactivated at 95 °C for 5 min was used as control.

Fixation was performed following 24 hrs of recombinant protein treatment using 4 % PFA / 4% Sucrose (Sigma) in 1x PBS at RT for 15 min. Neurons were subsequently washed 3 x with 1x PBS.

### Immunocytochemistry and immunohistochemistry

Tissue sections stained by mRNA FISH and fixed primary neurons were washed with 1 x PBS and blocked in 1 x PBS supplemented with 5 % (vol/vol) goat serum (Bio-Rad), 1 % (wt/vol) BSA (Sigma) and 0.1 % (vol/vol) Triton X-100 (Sigma) at RT for 1h. Primary antibodies were diluted in blocking solution and applied to samples at 4 °C overnight. Primary antibodies used in this study were: βAmyloid (BioLegend, 803001, RRID: AB_2564653, 1:200), GFAP (Sigma, G3893, RRID:AB_477010, 1:500), Homer (SynapticSystems, 160003, RRID:AB_887730, 1:500), Iba1 (Fujifilm Wako, 019-19741, RRID:AB_839504, 1:250), misfolded SOD1 (Médimabs, MM-0070-P, RRID:AB_10015296, 1:100), vGlut (MerckMillipore, AB5905, RRID:AB_2301751, 1:300); negative controls omitted the primary antibody. This was followed by 4 x 10 min washes in 1 x PBS at RT and subsequent application of suitable goat Alexa Fluor Plus secondary antibodies (488/546/647) diluted 1:500 in blocking solution at RT for 2 hrs. Samples were then washed 4 x 10 min with 1 x PBS at RT. Cryosections only were treated with 1x TrueBlack (Biotium) at RT for 30 sec to quench autofluorescence caused by the accumulation of lipofuscin and other protein aggregates, followed by 2 x washes with 1 x PBS. Nuclei of samples were counterstained with DAPI (Sigma; shown in blue in all confocal images) at 1 µg/ml in PBS and samples were mounted using DAKO Fluorescence Mounting Medium (Agilent).

### Microscopy

Stained tissue was imaged using a Zeiss LSM 880 confocal laser scanning microscope fitted with 40x (NA = 1.3) and 63x (NA = 1.4) objectives and photomultiplier tubes to detect fluorescence emission. For image acquisition, *xyz* confocal stacks were captured at a resolution of 1024 x 1024 pixels and at a step size of 1 µm. Microscope settings were established during first acquisition and subsequently not further modified. Four distinct fields of view were imaged from two representative sections per sample.

### Image processing and analysis

All images acquired from mouse tissue were processed and analysed in Fiji/ImageJ (Schindelin et al., 2012). *xyz* confocal stacks were collapsed into maximum *z* projections. Microgliosis was assessed by measuring both the number of microglia (DAPI^+^ nuclei embedded within typical microglial Iba1 immunoreactivity) and the total 2D surface area of Iba1 immunoreactivity within the acquired field of view. For area quantification, Iba1 immunoreactivity was processed by applying the ‘Remove Outliers’ function to remove non-specific noise (bright, radius = 2, threshold = 50), followed by thresholding at 35/255 to define the signal range, and two further rounds of the ‘Remove Outliers’ function, to fill-in nuclear and other gaps in Iba1 staining (dark, radius = 1), and to further remove non-specific noise (bright, radius = 3). The created Iba1 surface area was measured and used as a mask within which the *Dkk2* mRNA FISH signal surface area, thresholded to 30/255, was quantified.

Human frontal cortex image acquisitions were first subjected to ‘Linear unmixing’ with automatic fluorophore detection within the Zeiss Zen Black software (Zeiss) to remove overlapping signals between the five fluorophore channels. Unmixed and maximum z projected images were subsequently processed and analysed using the HALO FISH-IF v2.0.4 module (Indica Labs). The *DKK2* mRNA FISH signal surface area associated with *TREM2/P2RY12* double positive microglia cells was quantified. To achieve this, cell nuclei and their *xy* coordinates were recorded based on DAPI signal. Probe detection was optimised based on signal size, intensity of positive probe pixels and contrast threshold parameter settings (see Supplementary Table S1). The maximum distance threshold for probe signal assignment to nuclei was 25 µm. We classified cells positive for *P2RY12* and *TREM2* as microglia (DAPI^+^/*P2RY12/TREM2*^+^), determined their number, and measured the surface area of *DKK2* mRNA FISH signal associated with such DAPI^+^/*P2RY12/TREM2*^+^ cells.

Microglia-βAmyolid plaque distance analysis: we determined the 2D Euclidian distance of microglia to the proximal most βAmyolid plaque dense core in maximum projected images according to the following Fiji/ImageJ methodology: an intensity threshold was applied to the image channel containing βAmyloid immunostaining to identify the plaque dense core, which was usually more intensely labelled compared with the plaque periphery; due to the heterogeneous nature of βAmyloid plaques, threshold values were determined for each acquired image. The binary dense core image generated in the previous step was subjected to the ‘Exact Signed Euclidian Distance Transform (3D)’ (EDT) plug-in to create a 2D map where distance to the closest dense core was encoded in grey values from -1024 (furthest possible distance) to 0 (at dense core edge). *xy* position landmarks of DAPI^+^ microglia nuclear centres were placed on a binary image, which in turn was redirected to the EDT image in the ‘Set Measurements’ window, selecting ‘Mean grey value’ as measurement output. Note that *xy* positions of human microglia exported from HALO FISH-IF v2.0.4 module were imported into Fiji/ImageJ using the macro ‘ImportXYcoordinates.ijm’. Grey values at microglial *xy* positions were obtained using the ‘Analyze Particles’ function and converted into distance units by multiplying the grey value by the image *xy* pixel dimension (0.13495 µm) to yield microglia-βAmyloid plaque distances.

### Experimental design and statistical analysis

All means are stated ± standard deviation (SD). For the histological study aspects, *N* = number of subjects (humans or animals) and *n* = number of fields of view. For qualitative and quantitative histological assessments, we typically examined at least 4 subjects per group, imaging at least 4 different fields of view from 2 cryosections per subject, which met previously conducted sample size calculations according to Rosner (2015) with data inputs from Friedman et al., (2018). For the cytological study aspects, *N* = number of biological repeats, *n* = number of technical repeats (cells analysed). We used GraphPad Prism® software (GraphPad Software Inc.) for statistical analyses. D’Agostino and Pearson test was used to assess the normality of datasets. For the comparison of one independent variable between >2 groups, we used One-Way ANOVA with Tukey’s multiple comparison test. For statistical tests involving two independent variables we used Two-Way ANOVA with Šidák multiple comparisons test; where data points were missing, Mixed-effects analysis with Šidák multiple comparisons test was utilised. Significance was accepted at *p* ≤ 0.05.

### Data, software, and code availability

The data sets generated during and/or analysed during the current study are available from the corresponding authors on request. The Fiji/ImageJ macro ‘ImportXYcoordinates.ijm’ is available on the Github repository available via https://github.com/DominicAghaizu/ImageJMacros/blob/main/ImportXYcoordinates.ijm

## Results

### Microgliosis and microglial *Dkk2* upregulation in *APP^NL-G-F^* mice

We first investigated the microglial *Dkk2* expression pattern in the *APP^NL-G-F^* knock-in AD mouse model, which develops robust pathology from the physiological expression of humanised mouse amyloid precursor protein (*App*) harbouring Swedish, Beyreuther/Iberian, and Arctic mutations (Saito et al., 2014). To this end, we performed mRNA FISH on coronal brain cryosections to detect *Dkk2* mRNA *in situ* and acquired images from the motor cortex and the *stratum pyramidale*, with adjacent *stratum oriens* and *stratum radiatum*, of the hippocampal CA1 region. This was paired with immunohistochemical labelling using antibodies against Iba1 and βAmyloid to assess microglial *Dkk2* expression, as suggested previously (Friedman et al., 2018; Sala Frigerio et al., 2019; Meilandt et al., 2020), and to evaluate the spatial relationship between microglia and βAmyloid plaque lesions.

As expected, the brains of wild-type control littermate mice at 7 or 24 months were devoid of βAmyloid plaques and exhibited normally tiled Iba1^+^ microglia (Fig. 1*A,B*). In stark contrast, we detected βAmyloid plaques in the cortex and CA1 of transgenic *APP^NL-G-F^* mice at 7 and 24 months (Fig. 1*A,B*). This was accompanied by robust microgliosis as assessed by both normalised microglia cell count (DAPI^+^/Iba1^+^ cells) and area of Iba1 signal in maximum *z*- projected image stacks (Fig. 1*C;* note that the microglia spatial distribution will be addressed below). In the cortex, the number of microglia increased from 3.0 ± 0.2 to 6.5 ± 0.8 at 7 months and from 2.8 ± 0.8 to 15.9 ± 3.8 per field of view (FOV; equal to 1.8 x 10^-2^ mm^2^) at 24 months between control and *APP^NL-G-F^* mice; significant differences were found between timepoints and genotypes (Fig. 1*A,D,G*; Two-Way ANOVA, p = 0.0101 and p = 0.0002 respectively). Iba1 area increased from 821.9 ± 239.3 to 1505.8 ± 135.0 µm^2^ at 7 months and from 781.7 ± 121.1 to 1982.4 ± 471.6 µm^2^ at 24 months (Fig. 1A,*E*; Two-Way ANOVA, p = 0.1821 (timepoints) and p = 0.0018 (genotypes)). To assess microglial *Dkk2* expression levels, we quantified *Dkk2* mRNA FISH signal that was colocalised with Iba1 immunoreactivity (Fig. 1*C*). Comparing control and *APP^NL-G-F^* mice, the normalised area of *Dkk2* signal per DAPI^+^/Iba1^+^ microglial cell increased from 0.1 ± 0.1 to 0.3 ± 0.2 µm^2^ at 7 months and from 0.1 ± 0.1 to 1.2 ± 0.4 µm^2^ at 24 months (Fig. 1A,*F,G*; Two-Way ANOVA, p = 0.0245 (timepoints) and p = 0.0013 (genotypes)).

**Figure 1.**
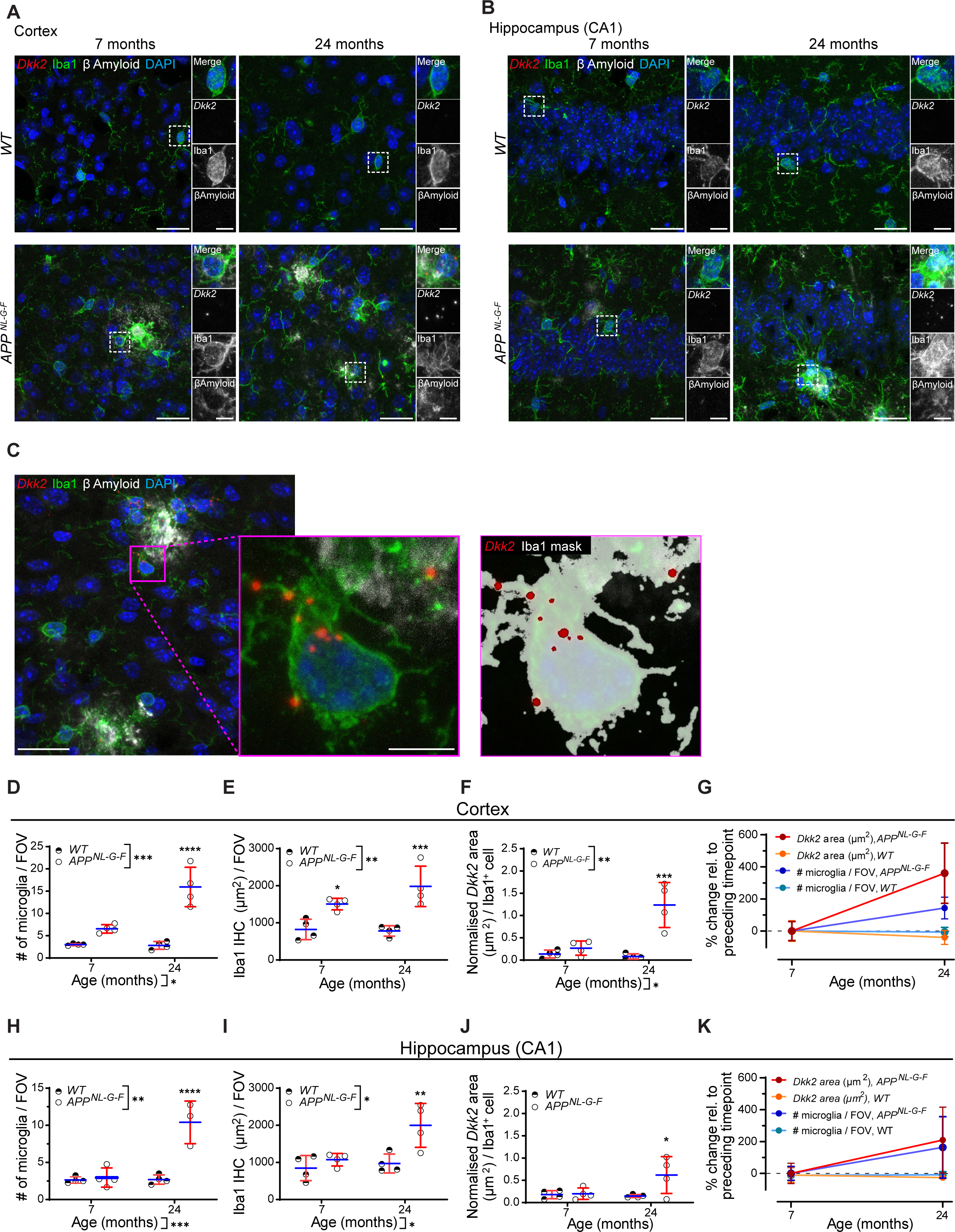
Microgliosis and microglial *Dkk2* upregulation in *APP^NL-G-F^* mice. *Dkk2* mRNA FISH as well as microglial Iba1 and βAmyloid IHC labelling in the motor cortex **(A)** and CA1 hippocampus **(B)** of *APP^NL-G-F^*mice. Boxed regions of interest (ROIs) were magnified for increased detail. **(C)** FIJI/ImageJ analysis workflow to quantifying punctated *Dkk2* mRNA FISH signal in Iba1-labelled (microglial) cells. Iba1 staining based analysis mask was generated, within which Dkk2 signal was quantified. **(D-G)** Microgliosis and *Dkk2* expression quantification in the *APP^NL-G-F^* motor cortex. **(D)** Quantification of microglia numbers per maximum projected field of view (FOV; 1.8 x 10^-2^ mm^2^). **(E)** Iba1 IHC surface area per maximum projected FOV. **(F)** Normalised *Dkk2* mRNA FISH signal area per DAPI^+^/Iba^+^ microglial cell. **(G)** Comparative % changes of *Dkk2* expression and microglia numbers during time course. **(H-K)** Microgliosis and Dkk2 expression quantification in the *APP^NL-G-F^* CA1 hippocampus. **(H)** Quantification of microglia numbers per maximum projected FOV. **(I)** Iba1 IHC surface area per maximum projected FOV. **(J)** Normalised *Dkk2* mRNA FISH signal area per DAPI^+^/Iba^+^ microglial cell. **(K)** Comparative % changes of *Dkk2* expression and microglia numbers during time course. Individual data points represent the average of 4 FOVs analysed for each animal **(D-F, H-J)** or total averages from all animals per group **(G, K)**. N = 4 animals per condition and time point, n = 4 different fields of view / animal and brain region. Scale bars, (A-C) 25 µm; magnified ROIs: 5 µm. Two-way ANOVA with Multiple comparisons test. *p < 0.05; **p < 0.01; ***p < 0.001; ****p < 0.0001.

Similar patterns of microgliosis and *Dkk2* upregulation were observed in the hippocampal CA1 region. The number of microglia increased from 2.6 ± 0.4 to 3.0 ± 1.1 (n.s.) at 7 months and from 2.7 ± 0.5 to 10.4 ± 2.3 at 24 months between control and *APP^NL-G-F^* mice (Fig. 1B,*H,K*; Mixed-effects analysis, p = 0.0005 (timepoints) and p = 0.0051 (genotypes)). Iba1 area increased from 846.9 ± 292.8 to 1072.6 ± 146.3 µm^2^ at 7 months and from 970.6 ± 221.3 to 1997.7 ± 511.8 µm^2^ at 24 months (Fig. 1*B*,*I*; Two-Way ANOVA, p = 0.0119 (timepoints) and p = 0.0288 (genotypes)). *Dkk2* expression per microglial cell quantified by mRNA FISH remained unchanged at 7 months (0.2 ± 0.1 to 0.2 ± 0.1 µm^2^) but rose from 0.1 ± 0.0 to 0.6 ± 0.4 µm^2^ at 24 months (Fig. 1B,*J,K*; Two-Way ANOVA, p = 0.1363 (timepoints) and p = 0.0652 (genotypes)).

Taken together, our data demonstrate robust microgliosis in conjunction with *Dkk2* upregulation in *APP^NL-G-F^*mice compared with littermate controls, adding a spatial dimension to a previously published single-cell RNA sequencing (RNA-Seq) study that identified *Dkk2* expression in DAM/ARM microglia of the same mouse model (Sala Frigerio et al., 2019).

### Microgliosis and microglial *Dkk2* upregulation in *APP/PS1* mice

Following investigation of *APP^NL-G-F^* mice, we assessed microgliosis and *Dkk2* upregulation in a second AD mouse model, the *APP/PS1* mouse, that expresses chimeric mutant mouse/human *App* and mutant human *presenilin 1*, both associated with early onset familial AD in humans (Jankowsky et al., 2004).

βAmyloid plaque load progressively increased in *APP/PS1* mice starting from 8 months, whereas wild-type control littermates lacked βAmyloid plaques altogether. This was especially evident in the cortex (Fig. 2*A*). While plaques were detectable in the hippocampus of *APP/PS1* mice (data not shown), CA1 *stratum pyramidale* proximal regions – the standardised hippocampal brain region that was imaged in our study – rarely exhibited plaque depositions (Fig. 2*B*).

**Figure 2.**
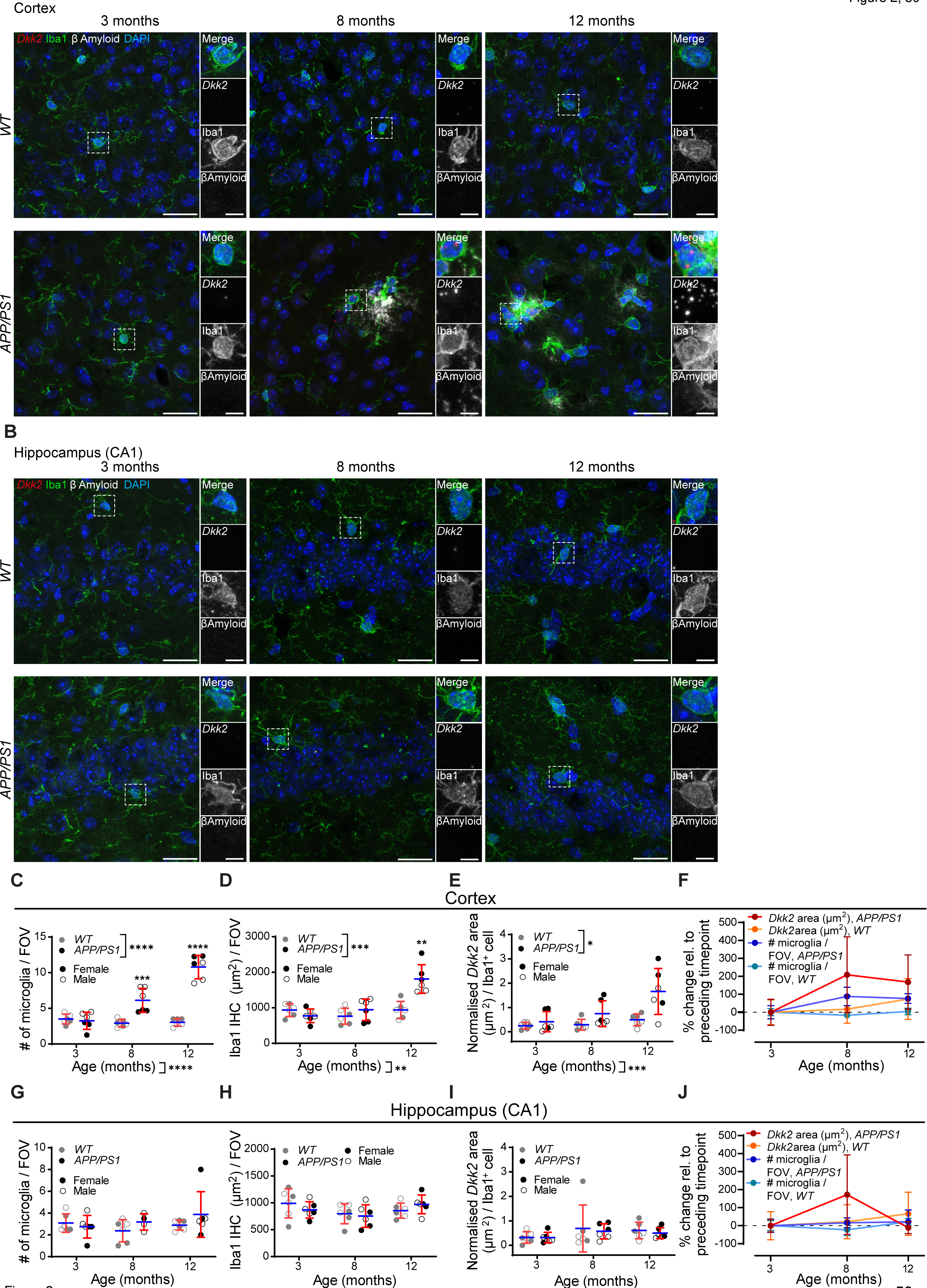
Microgliosis and microglial *Dkk2* upregulation in *APP/PS1* mice. *Dkk2* mRNA FISH as well as microglial Iba1 and βAmyloid IHC labelling in the motor cortex **(A)** and CA1 hippocampus **(B)** of *APP/PS1* mice. Boxed regions of interest (ROIs) were magnified for increased detail. **(C-F)** Microgliosis and *Dkk2* expression quantification in the *APP/PS1* motor cortex. **(C)** Quantification of microglia numbers per maximum projected FOV (FOV = 1.8 x 10^-2^ mm^2^). **(D)** Iba1 IHC surface area per maximum projected FOV. **(E)** Normalised *Dkk2* mRNA FISH signal area per DAPI^+^/Iba^+^ microglial cell. **(F)** Comparative % changes of *Dkk2* expression and microglia numbers during time course. **(G-J)** Microgliosis and Dkk2 expression quantification in the *APP/PS1* CA1 hippocampus. **(G)** Quantification of microglia numbers per maximum projected FOV. **(H)** Iba1 IHC surface area per maximum projected FOV. **(I)** Normalised *Dkk2* mRNA FISH signal area per DAPI^+^/Iba^+^ microglial cell. **(K)** Comparative % changes of *Dkk2* expression and microglia numbers during time course. Individual data points represent the average of 4 FOVs analysed for each animal **(C-E, G-I)** or total averages from all animals per group **(F, J)**. N = 6 animals (3x females, 3x males) per time point and condition, n = 4 different fields of view / animal and brain region. Scale bars, (A-C) 25 µm; magnified ROIs: 5 µm. Two-way ANOVA with Multiple comparisons test. *p < 0.05; **p < 0.01; ***p < 0.001; ****p < 0.0001.

While the number of DAPI^+^/Iba1^+^ microglia remained unchanged in control versus *APP/PS1* mice at 3 months (3.5 ± 0.6 to 3.3 ± 1.1 per FOV), it significantly increased in *APP/PS1* mice at 8 months (2.9 ± 0.5 to 6.1 ± 1.5) and at 12 months (3.0 ± 0.5 to 10.8 ± 1.5) (Fig. 2*A*,*C*; Two-Way ANOVA, p < 0.0001 (timepoints) and p < 0.0001 (genotypes)). Iba1 area did not markedly differ at 3 months (935.4 ± 164.0 vs 774.8 ± 175.2 µm^2^) and at 8 months (764.1 ± 205.3 vs 945.1 ± 275.8 µm^2^) but was significantly increased in *APP/PS1* mice at 12 months (937.9 ± 220.8 vs 1811.1 ± 367.9 µm^2^) (Fig. 2*A*,*D*; Two-Way ANOVA, p = 0.0030 (timepoints) and p = 0.0006 (genotypes)). Microgliosis was accompanied by progressively increasing *Dkk2* expression per microglial cell at the mRNA level: 0.9 ± 0.6 to 1.2 ± 0.9 µm^2^ at 3 months, 1.0 ± 0.7 to 3.6 ± 2.5 µm^2^ at 8 months, and 1.8 ± 1.2 to 9.7 ± 5.5 µm^2^ at 12 months (Fig. 2*A*,*E*; Two-Way ANOVA, p = 0.0003 (timepoints) and p = 0.0391 (genotypes)). The rate of increase of microgliosis (number of microglia) and *Dkk2* expression in *APP/PS1* mice was rapid between the ages of 3 and 8 months (88.3 ± 46.9 % for microgliosis, 208.6 ± 192.9 5 for *Dkk2* expression), at which point it plateaued (76.3 ± 24.6 % for microgliosis, 167.9 ± 138.2 % for *Dkk2* expression) (Fig. 2*A,F*). In agreement with published literature (Wang et al., 2003), we further noted that for the quantified metrics described above, female mice usually exhibited a more severe phenotype, especially at the final 12 months time point (Fig. 2*C,D,E*).

As noted above, hippocampal CA1 *stratum pyramidale* proximal regions in *APP/PS1* mice were mostly devoid of βAmyloid plaques. Here, we were unable to detect any changes in the number of DAPI^+^/Iba1^+^ microglia (Fig. 2*B,G,J*), Iba1 area (Fig. 2*B,H*), and *Dkk2* mRNA signal per microglial cell compared with littermate controls (Fig. 2*B,I,J*) (Two-Way ANOVA, all n.s.).

Thus, we were able to largely replicate our findings regarding microgliosis and microglial *Dkk2* upregulation in two widely used AD mouse models (*APP^NL-G-F^*and *APP/PS1* mice), again adding spatial information to a previously published meta-analysis of single-cell RNA-Seq datasets (Friedman et al., 2018). However, the lack of βAmyloid plaques and microglial phenotype in hippocampal CA1 *stratum pyramidale* proximal regions of the *APP/PS1* mouse evokes the notion that the microglial phenotype investigated here could be linked to plaque proximity.

### *Dkk2^+^* microglia exhibit increased propensity for clustering around βAmyloid plaques

To investigate whether *Dkk2* expression status was correlated with βAmyloid plaque proximity, we performed nearest neighbour analysis to quantify the spatial relationship between microglia and the nearest βAmyloid plaque dense core identified following βAmyloid IHC in *APP^NL-G-F^* and *APP/PS1* mice (schematic shown in Fig. 3*A*). Frequency distributions of recorded distances were summarised in histograms. In AD mouse models, we distinguished between *Dkk2^+^* and *Dkk2^-^* microglia, whereas no such distinction was made in wild-type mice as *Dkk2* expression levels were negligible at all time points (Fig. 1*A*,*B*,*F*,*J*, 2*A*,*B*,*E*,*I*). Furthermore, where no plaques were evident (e.g., in wild-type or pre-disease stage mice or in some hippocampal CA1 *stratum pyramidale* proximal regions) distances to plaque dense core “placeholders” randomly placed on confocal images were measured instead.

**Figure 3.**
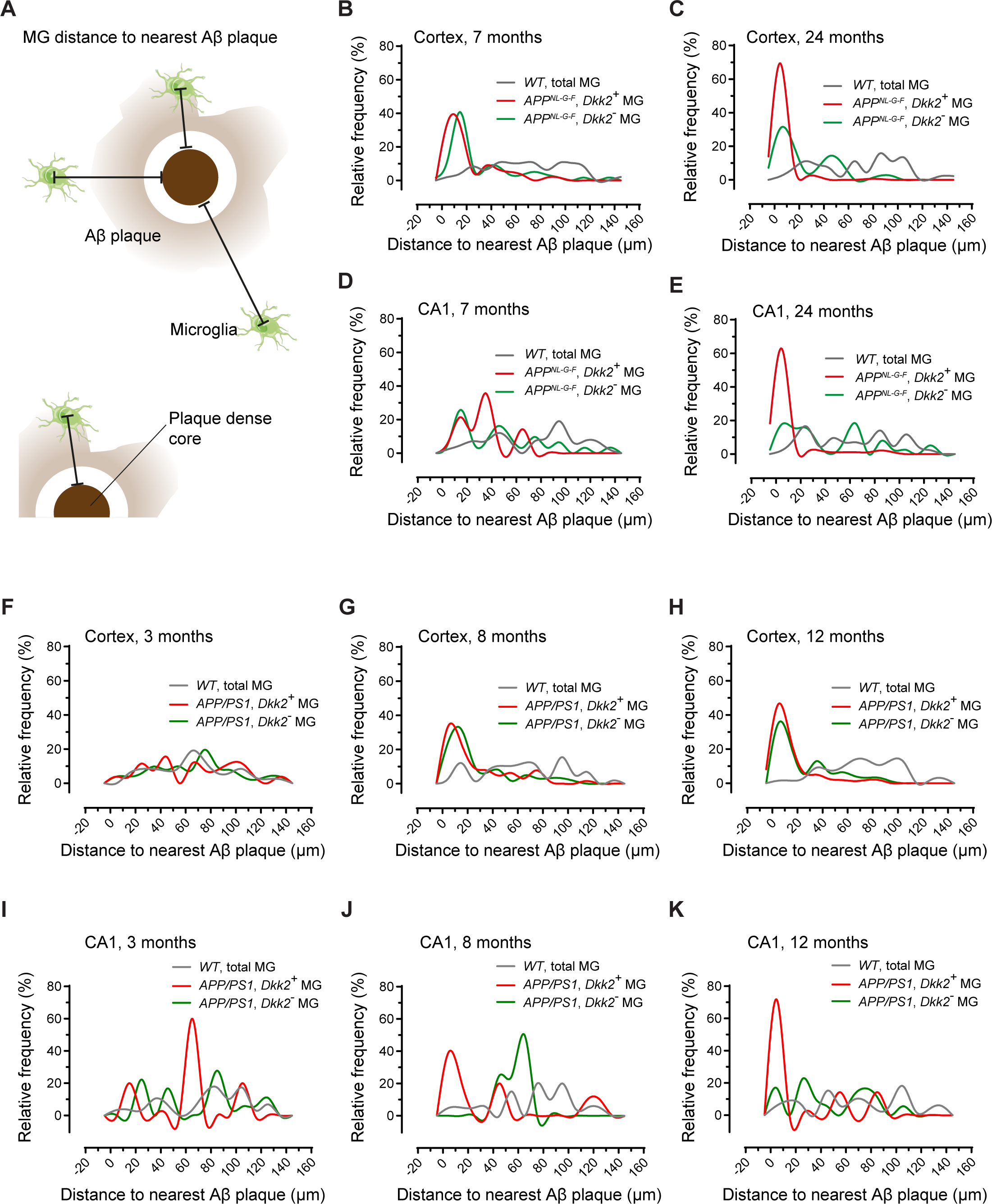
*Dkk2^+^* microglia cluster around βAmyloid plaques in *APP^NL-G-F^* and *APP/PS1* mice. (A) Schematic showing methodology of measuring distances between microglia and nearest βAmyloid plaque dense core. **(B-E)** Distribution of microglia (*Dkk2^+^*, *Dkk2^-^* or total microglia (MG) populations) distances to nearest βAmyloid plaque dense core in *APP^NL-G-^*^F^ or littermate control mice. Relative frequency distribution in the *APP^NL-G-^*^F^ or control motor cortex at 7 months **(B)** and 24 months **(C)**. Relative frequency distribution in the *APP^NL-G-^*^F^ or control CA1 hippocampus at 7 months **(B)** and 24 months **(C)**. **(F-K)** Distribution of microglia (*Dkk2^+^*, *Dkk2^-^*or total microglia (MG) populations) distances to nearest βAmyloid plaque dense core in *APP/PS1* or littermate control mice. Relative frequency distribution in the *APP/PS1* or control motor cortex at 3 months **(F)**, 8 months **(G)**, and 12 months **(H)**. Relative frequency distribution in the *APP/PS1* or control CA1 hippocampus at 3 months **(I)**, 8 months **(J)**, and 12 months **(K)**. *APP^NL-G-^*^F^/control: N = 4 animals per condition and time point, n = 4 different fields of view / animal and brain region; *APP/PS1*/control: N = 6 animals (3x females, 3x males) per time point and condition, n = 4 different fields of view / animal and brain region.

As would be expected, wild-type littermate controls of *APP^NL-G-F^* mice used in our study exhibited microglia at varying/random distances to the nearest randomly assigned dense core placeholder in the motor cortex and CA1 hippocampus at 7 and 24 months (Fig. 1*A,B*, 3*B-E*). This finding is in keeping with the homogeneous tiling behaviour usually exhibited by microglia in the healthy CNS (Nimmerjahn et al., 2005). In stark contrast, a large proportion of *Dkk2^+^* and *Dkk2^-^* microglia were found within 20 µm of the nearest plaque dense core in the cortex of 7 months old *APP^NL-G-F^* mice, while *Dkk2^+^* microglia were predominantly located within 40 µm of plaque dense cores in the CA1 hippocampus (Fig. 3*B,D*). By 24 months, the clustering of microglia around βAmyloid plaque dense cores, especially that of *Dkk2^+^* microglia, became even more pronounced both in the cortex and in the CA1 hippocampus (Fig. 3*C,D*).

We observed similar plaque-microglia distance relationships in *APP/PS1* mice and their respective wild-type littermate controls. Microglia in the wild-type mouse cortex and CA1 hippocampus were evenly distributed relative to the nearest randomly placed plaque dense core placeholder (Fig. 3*F-K*). In the cortex of 3 months old (pre disease stage and plaque free) *APP/PS1* mice, both *Dkk2^+^* and *Dkk2^-^*microglia exhibited similar wild-type-like distance distributions (Fig. 3*F*), whereas microglia increasingly clustered within 20 µm of plaque dense cores at subsequent (disease stage) time points, with *Dkk2^+^* microglia exhibiting slightly more pronounced clustering versus *Dkk2^-^* microglia at 12 months (Fig. 3*G,H*). As discussed above, due to the small amounts of βAmyloid plaques in hippocampal CA1 *stratum pyramidale* proximal regions of the *APP/PS1* mouse, microglia distributions were comparatively variable, especially at 3 months (Fig. 3*I*), even though substantial clustering of *Dkk2^+^* microglia was registered in those instances were βAmyloid plaques were observed in CA1 *stratum pyramidale* proximal regions at 8 and 12 months (Fig. 3*J,K*).

While it is widely known that microglia accumulate around CNS lesions such as βAmyloid plaques, our data further suggest that clustering around plaques is frequently accompanied by the expression of *Dkk2*. Conversely, in the healthy brain, microglia were evenly tiled and lacked *Dkk2* expression.

### Microgliosis and microglial *Dkk2* upregulation in *SOD1^G93A^* AL*S* mice

Having demonstrated microgliosis and clustering of *Dkk2^+^* microglia around βAmyloid plaques in two different widely used AD mouse models, we next investigated whether our findings could be recapitulated in another neurodegeneration mouse model, the *SOD1^G93A^* amyotrophic lateral sclerosis (ALS) mouse (Gurney et al., 1994). According to the meta-analysis of single-cell RNA-Seq datasets by Friedman et al., 2018, microglial *Dkk2* upregulation should also be evident in this mouse model. It expresses the mutant human *SOD1^G93A^* gene that causes motor neuron degeneration in the spinal cord and other parts of the CNS, which underlies ALS (Gurney et al., 1994). We performed mRNA FISH to detect microglial *Dkk2* mRNA *in situ* paired with immunohistochemical labelling using an antibody against Iba1 on transverse cryosections from the lumbar (L)5 region of the spinal cord and acquired images from the ventral horn, an area that displays robust motor neuron degeneration in this mouse model (Gurney et al., 1994), In control mice expressing wild-type human *SOD1 (SOD1^WT^)*, but not in mice expressing *SOD1^G93A^,* no overt changes in microgliosis and *Dkk2* expression were observed at any of the assessed time points (Fig. 1*A-E*). At the early 50 days time point, *SOD1^G93A^* mice still exhibited control levels of microgliosis (1.7 ± 0.1 versus 1.6 ± 0.2 DAPI^+^/Iba1^+^ microglia per FOV (Fig. 4*A*,*B*), 244.7 ± 73.2 vs 266.5 ± 59.7 µm^2^ Iba1 area (Fig. 4*A*,*C*)). However, the number of microglia significantly rose from 1.8 ± 0.2 to 6.3 ± 0.9 at 100 days and from 1.5 ± 0.4 to 13.6 ± 0.9 at 120 days (Fig. 4*A*,*B*; Two-Way ANOVA, p < 0.0001 (timepoints) and p = 0.0001 (genotypes)). Accordingly, the area of Iba1 immunoreactivity increased from 328.1 ± 75.3 to 918.8 ± 31.7 µm^2^ at 100 days and from 248.7 ± 42.1 to 1646.0. ± 184.7 µm^2^ at 120 days (Fig. 4*A*,*C*; Two-Way ANOVA, p < 0.0001 (timepoints) and p = 0.0001 (genotypes)). *Dkk2* expression per microglial cell progressively increased in *SOD1^G93A^* versus *SOD1^WT^* mice, although this increase only reached significance at 120 days: 0.7 ± 0.5 vs 0.8 ± 0.7 µm^2^ at 50 days, 0.5 ± 0.3 vs 2.4 ± 0.8 µm^2^ at 100 days, and 0.5 ± 0.6 vs 10.7 ± 4.6 µm^2^ at 120 days (Fig. 4*A*,*D*; Two-Way ANOVA, p = 0.0154 (timepoints) and p = 0.0193 (genotypes)). Thus, fast-paced microgliosis is evident between 50 and 100 days in *SOD1^G93A^* mice, with a slightly reduced rate of acceleration between 100 days and 120 days (Fig. 4*E*). Conversely, microglial *Dkk2* upregulation appears to accelerate especially in the final pathological stages.

**Figure 4.**
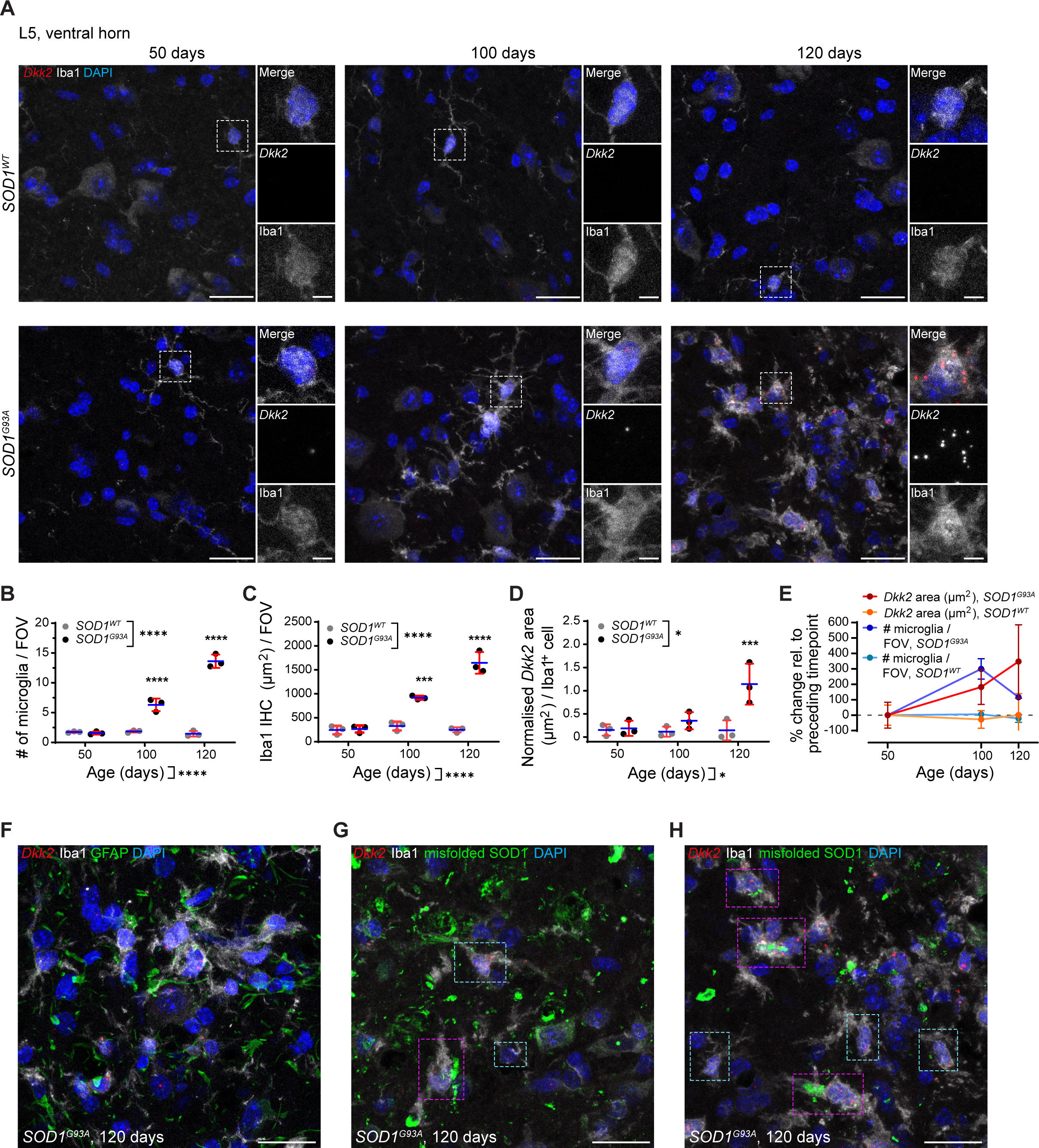
Microgliosis and microglial *Dkk2* upregulation in *SOD1^G93A^* AL*S* mice. (A) *Dkk2* mRNA FISH and microglial Iba1 IHC labelling in the L5 spinal cord ventral horn of mice transgenically expressing human *SOD1^WT^* (control) or mutant *SOD1^G93A^* at 50, 100, and 120 days. Boxed regions of interest (ROIs) were magnified for increased detail. **(B)** Quantification of microglia numbers per maximum projected FOV (FOV = 1.8 x 10^-2^ mm^2^). **(C)** Iba1 IHC surface area per maximum projected FOV. **(D)** Normalised *Dkk2* mRNA FISH signal area per DAPI^+^/Iba^+^ microglial cell. **(E)** Comparative % changes of *Dkk2* expression and microglia numbers during time course. **(F)** *Dkk2* mRNA FISH together with microglial Iba1 and astroglial GFAP IHC labelling in the L5 spinal cord ventral horn of 120 days old *SOD1^G93A^* mice. **(G,H)** *Dkk2* mRNA FISH together with microglial Iba1 and misfolded SOD1 IHC labelling in the L5 spinal cord ventral horn of 120 days old *SOD1^G93A^* mice. Magenta and cyan ROIs respectively depict proximity and absence of clear association between DAPI^+^/Iba1^+^ microglia and misfolded SOD1 foci. Individual data points represent the average of 4 FOVs analysed for each animal **(C-D)** or total averages from all animals per group **(E)**. N = 3 animals per time point and condition, n = 4 fields of view per animal. Scale bars, (A,F-H) 25 µm; magnified ROIs: 5 µm. Two-way ANOVA with Multiple comparisons test. *p < 0.05; **p < 0.01; ***p < 0.001; ****p < 0.0001.

We next sought to investigate whether microgliosis and microglial *Dkk2* upregulation in the *SOD1^G93A^*ALS mouse model were spatially correlated with local CNS lesions, analogous to that observed in the *APP^NL-G-F^* and *APP/PS1* AD mouse models. In absence of AD-typical βAmyloid plaques in ALS, we combined *Dkk2* mRNA FISH and microglial immunohistochemical labelling with the immunolabelling of GFAP to visualise astrocytes and immunolabelling of misfolded SOD1 to visualise aggregates of misfolded mutant SOD1^G93A^. In 120 days old *SOD1^G93A^* ALS mice, we failed to detect clustering of microglia, irrespective of their Dkk2 expression status, specifically around GFAP (Fig. 4*F*). However, we observed some degree of microglial clustering around misfolded SOD1 immunoreactivity (Fig. 4*G,H;* magenta regions of interest (ROIs)). However, many microglia did not exhibit local accumulation around misfolded SOD1 lesions (Fig. 4*G,H*: cyan ROIs). In absence of a clear clustering pattern, these observations were not quantified.

Taken together, the microgliosis and microglial *Dkk2* upregulation detected in the brains of AD mouse models could also be replicated in an unrelated neurodegeneration mouse model, namely in the spinal cord of the *SOD1^G93A^* ALS mice. While some degree of clustering around misfolded SOD1 aggregates occurred, this was not as robust as the clustering around βAmyloid plaques in the *APP^NL-G-F^* and *APP/PS1* AD mouse models. Nonetheless, our findings support the published notion that *Dkk2* upregulation may be part of a general response in CNS microglia as they transition from surveillance to activation (DAM/ARM microglia), at least in mouse models of neurodegeneration (Friedman et al., 2018; Sala Frigerio et al., 2019; Meilandt et al., 2020). This supports the possibility that *Dkk2* represents a DAM/ARM marker gene, at least in mice.

### DKK2 recombinant protein disrupts WNT7a-induced synapse features in cultured neurons

We next sought to investigate what effect Dkk2 protein secreted by microglia might have on its surroundings. We focused our study on synapses in mature primary neuron cultures due to the well-known anti-synaptic effect that the Dkk2 homolog Dkk1 has on them, which it brings about by decreasing canonical and increasing non-canonical Wnt signalling (Purro et al., 2012; Galli et al., 2014; Marzo et al., 2016; Elliott et al., 2018; Sellers et al., 2018).

To this end, we treated mature rat hippocampal neuron cultures sparsely expressing *hSyn:EGFP* at 21 days *in vitro* with recombinant proteins for 24 hrs (WNT7a, 200 ng/ml; DKK1, 100 ng/ml; DKK2, 100 ng/ml; DKK2 + WNT7a, 100 and 200 ng/ml; BSA control, 100 ng/ml). This was followed by immunocytochemical labelling using antibodies against the pre- and post-synaptic markers vGlut and Homer. A typical sparsely labelled (*hsyn:EGFP^+^*) neuron with highlighted primary dendrite (boxed ROI) that was used for analysis is depicted in Fig. 5*A*. WNT7a treatment significantly increased the number of dendritic spines as well as the number of post-synaptic homer puncta compared with BSA treatment (Fig. 5*B,C,G,H;* One-Way ANOVA, dendritic spines: p = 0.0023; homer puncta: p = 0.0309). Conversely, these metrics were unaffected by DKK1 and DKK2 treatment (Fig. 5*D,E,G,H;* One-Way ANOVA; all n.s.) and crucially also by combined DKK2 + WNT7a treatment (Fig. 5*F,G,H;* One-Way ANOVA; n.s.). The absolute number of synapses (defined as Homer/vGlut apposition events with up to 1 µm distance) was similarly increased by WNT7a but not by DKK1, DKK2 or a combination of DKK2 and WNT7a compared with BSA, although this did not reach statistical significance (Fig. 5*I*; One-Way ANOVA; n.s.).

**Figure 5.**
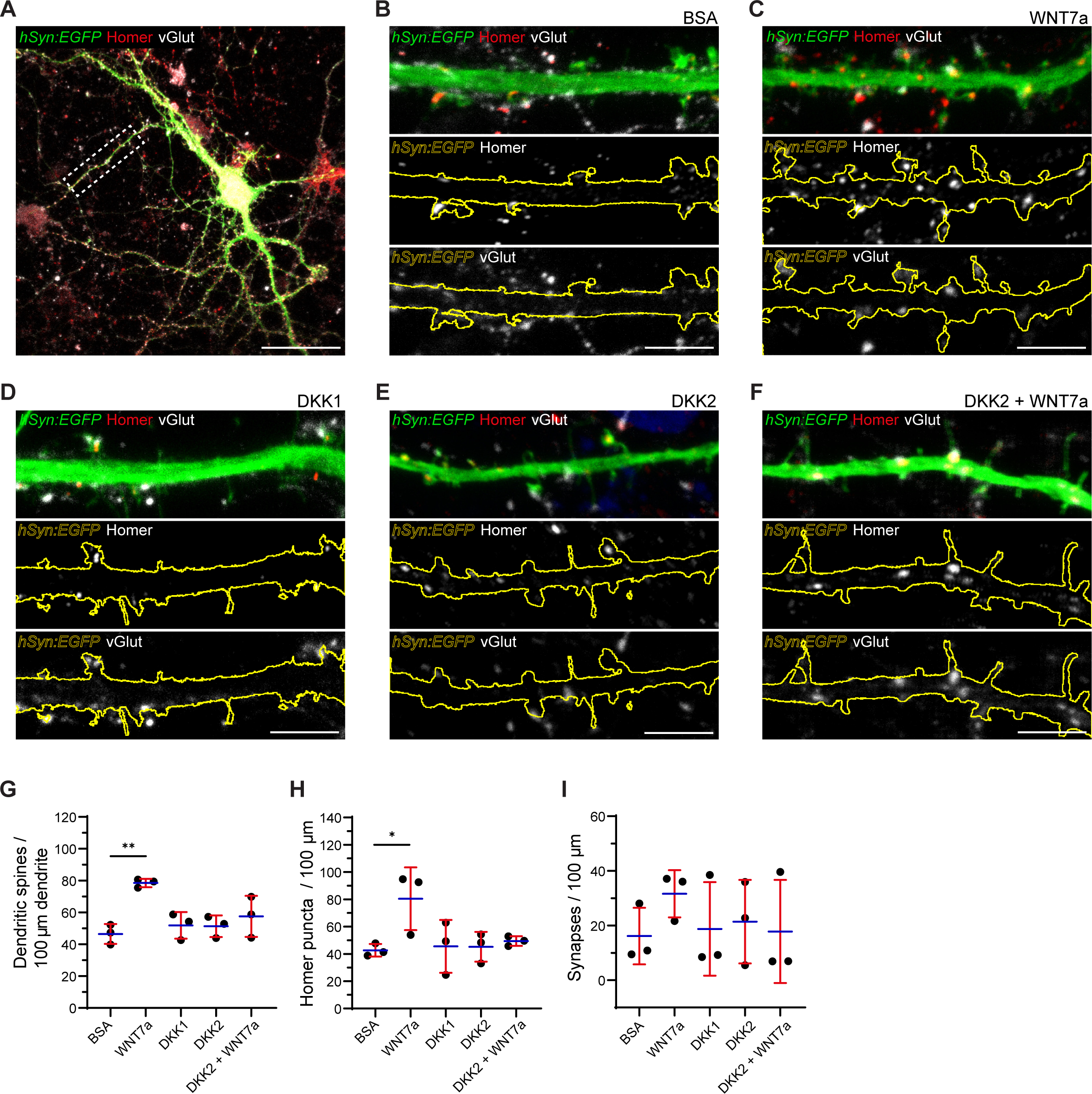
Recombinant DKK2 protein neutralises the synaptogenic effect of WNT7a in mature hippocampal primary neurons. (A) Typical rat hippocampal neuron at DIV22 expressing *hSyn:EGFP* immunolabelled with Homer and vGlut. Boxed ROI indicates a primary dendritic branch, on which analysis in this section was focused. **(B-F)** Representative primary dendrites of DIV22 hippocampal neurons treated for 24 hrs with 100 ng/ml BSA control **(B)**, 200 ng/ml WNT7a **(C)**, 100 ng/ml DKK1 **(D)**, 100 ng/ml DKK2 **(E)**, and 100 ng/ml / 200 ng/ml DKK2 + WNT7a **(F)**. Immunolabelling for the pre- and post-synaptic markers vGlut and homer was performed, and merged views are shown in top panels. Remaining panels show homer (middle panel) and vGlut (lower panel) with outlined primary dendrite boundaries based on *hSyn:EGFP* labelling. **(G)** Normalised number of dendritic spines per 100 µm primary dendrite. **(H)** Normalised number of homer puncta per 100 µm primary dendrite. **(I)** Normalised number of synapses (defined as homer/vGlut apposition events with a maximum distance of 1 µm) per 100 µm primary dendrite. **(J)** Relative number of homer puncta apposed within 1 µm by vGlut. N = 3 biological repeats, n = at least 15 analysed neurons per condition. Scale bars, (A) 50 µm; (B-F) 5 µm. One-way ANOVA with Multiple comparisons test. *p < 0.05; **p < 0.01.

We thus infer from our data that WNT7a signalling tone was low/absent in our rat hippocampal primary neuron cultures (DKK1 and DKK2 did not reduce synaptic metrics to levels below those found with BSA treatment). Crucially however, DKK2 treatment completely abolished the pro-synaptogenic effects of WNT7a treatment.

### *DKK2* is not upregulated in human microglia

We have thus far demonstrated significant microgliosis and microglial *Dkk2* upregulation in AD and ALS mouse models of neurodegeneration, as well as clustering of *Dkk2^+^* microglia around βAmyloid plaques. In combination with previously published studies, which have demonstrated microglial *Dkk2* upregulation by single-cell RNA-Seq (Friedman et al., 2018; Sala Frigerio et al., 2019; Meilandt et al., 2020), this led us to postulate that *Dkk2* may represent a *bona fide* DAM/ARM marker gene at least in neurodegeneration mouse models. We next sought to investigate whether our findings were recapitulated in human subjects diagnosed with AD.

To analyse microglial *DKK2* expression in humans, we obtained human *post-mortem* frontal cortex brain tissue from healthy control individuals, as well as individuals diagnosed with AD and pathological ageing, the latter being defined as non-demented individuals with AD-typical histopathologic changes. Demographic data and post-mortem brain assessments are summarised in Supplementary Table S2. We performed mRNA FISH to detect *DKK2* mRNA in microglia that were additionally labelled by mRNA FISH for the microglial markers *TREM2* and *P2RY12*. This was paired with immunohistochemical labelling using an antibody against βAmyloid to label βAmyloid plaques. As expected, samples from control individuals were devoid of βAmyloid plaques while those classified “pathological ageing” and “AD” exhibited progressively increasing levels of plaque burden (Fig. 6*A-C*). However, we did not detect significant differences in the number of DAPI^+^/*TREM2^+^*/*P2RY12^+^*microglia per field of view between control, pathological ageing, and AD groups (Fig. 6*A-D*; control: 11.1 ± 10.7 microglia

**Figure 6.**
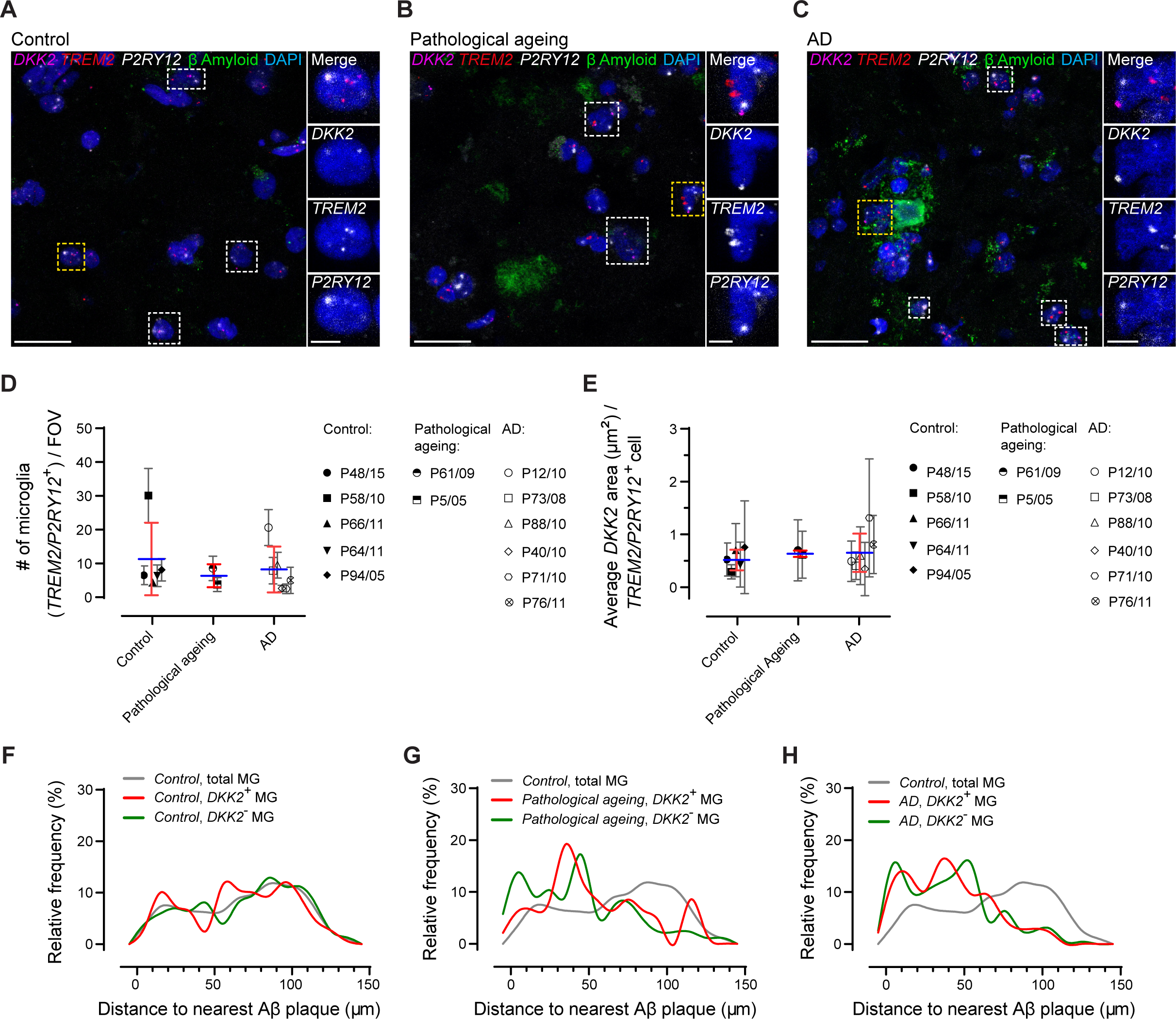
*DKK2* is *not* upregulated at the mRNA level in post-mortem brains from AD patients. (A-C) Representative confocal images depicting microglial *DKK2* expression in the human frontal cortex. *DKK2* as well as microglial *TREM2* and *P2RY12* mRNA FISH signal in conjunction with βAmyloid IHC labelling in post-mortem human frontal cortex samples from healthy control individuals **(A)**, individuals diagnosed with pathological ageing **(B)**, and individuals diagnosed with AD **(C)**. Boxed ROIs highlight microglia expressing *DKK2* (DAPI^+^/*DKK2^+^*/*TREM2^+^*/*P2RY12^+^*); yellow boxed ROIs were enlarged for improved visualisation. **(D)** Quantification of microglia (DAPI^+^/*TREM2^+^*/*P2RY12^+^*) numbers per maximum projected FOV (FOV = 1.8 x 10^-2^ mm^2^). **(E)** Normalised *DKK2* mRNA FISH signal area per DAPI^+^/*TREM2^+^*/*P2RY12^+^*microglial cell. **(F-H)** Distribution of DAPI^+^/*TREM2^+^*/*P2RY12^+^*microglia (*Dkk2^+^*, *Dkk2^-^* or total microglia (MG) populations) distances to nearest βAmyloid plaque dense core in post-mortem human frontal cortex samples. Individual plots show relative frequency distributions in individuals classified as healthy control **(F)**, pathological ageing **(G)**, and AD **(H)**. Healthy control individuals: N = 5 individuals, n = 8 fields of view); AD (Braak & Braak stage 5-6): N = 6 individuals, n = 8 fields of view; pathological ageing (Braak & Braak stage 3-4): N = 2 individuals, n = 8 fields of view. Data points represent the average of 4 FOVs analysed for each individual subject (mean ± SD); individual subject mean values were further averaged for each group of interest and summarised as mean ± SD (blue horizontal bars, red error bars). One-way ANOVA with Multiple comparisons test. No statistical differences identified. Scale bars, (A-C) 25 µm; (A-C enlarged ROIs) 5 µm. See also Supplementary Table T1.

/ FOV; pathological ageing: 6.2 ± 3.4; AD: 8.0 ± 6.8; one-way ANOVA, p = 0.4507). Similarly, *DKK2* expression per DAPI^+^/*TREM2^+^*/*P2RY12^+^* microglial cell did not differ between control (0.5 ± 0.2 µm^2^), pathological ageing (0.7 ± 0.1 µm^2^), and AD groups (0.7 ± 0.4 µm^2^) (Fig. 6*A-C,E;* one-way ANOVA, p = 0.7689).

We subsequently assessed the clustering behaviour of microglia around βAmyloid plaques. In healthy control individuals, the total microglia population displayed a varying/random spatial distribution around the nearest randomly placed dense core placeholder, which furthermore did not appear to be modified by *DKK2* expression status (Fig. 6*A,F*). In individuals classified as “pathological ageing”, we identified emerging populations of both *DKK2^+^* and *DKK2^-^* microglia that frequently accumulated around βAmyloid plaque dense cores up to a distance of 50 µm, although clustering in the proximal most regions was more robust for *DKK2^-^*cells (Fig. 6*B,G*). This clustering was further consolidated, especially among *DKK2^+^*microglia, whose predominant distribution now also included proximal most regions (Fig. 6*C,H*).

Taken together, our data on human frontal cortex post-mortem tissue indicate that neither the increase in microglial numbers nor microglial *DKK2* upregulation, both of which were evident in mouse models, occur in human brains under conditions classified as “pathological ageing” and “AD”. However, microglia did exhibit clustering behaviour around βAmyloid plaques even though this was not linked to *DKK2* expression.

## Discussion

Past and present research have linked dysregulated Wnt signalling to AD (e.g. Palomer et al., 2022; previously reviewed by Purro et al., 2014; Palomer et al., 2019; Aghaizu et al., 2020; Inestrosa et al., 2021). However, recent research has also more intimately linked microglia and neuroinflammation to AD, as initially exemplified by variants of genes predominantly expressed in microglia like *TREM2* and *CD33* exhibiting disease modifying properties (Bradshaw et al., 2013; Guerreiro et al., 2013). New evidence even suggests that the microglial AD response is itself regulated by Wnt signalling, as the signalling pathway downstream of TREM2, essential for regulating microglial survival and proliferation, cross-talks with the Wnt pathway (Zheng et al., 2017; Meilandt et al., 2020).

Here we sought to explore the role of *DKK2/Dkk2*, which encodes a Wnt signalling modulator, that was upregulated in a subpopulation of microglia (DAM/ARM) in various single and bulk cell RNA-Seq studies on neurodegeneration mouse models (Friedman et al., 2018; Sala Frigerio et al., 2019; Meilandt et al., 2020). Our histological data obtained largely by mRNA FISH combined with immunocytochemistry replicated the findings cited above. Crucially however, we added valuable spatial information on the location of *Dkk2^+^* microglia with respect to neurodegenerative lesions such as βAmyloid plaques in AD mouse models, where *Dkk2^+^*microglia exhibited a potential to cluster near βAmyloid plaques that was greater or at least equal to that of *Dkk2^-^* microglia. The exact role of Dkk2 protein expression is yet to be fully understood, but assuming its reported role as a secreted, soluble protein (reviewed by Niehrs, 2006) we speculated that Dkk2’s mechanism of action could be autocrine or paracrine in nature. In support of the former, oncological evidence suggests that peripheral immune natural killer and CD8^+^ T cells, which are derived from the same myeloid lineage as CNS microglia, can detect soluble Dkk2. However, in this context, Dkk2 was utilised as an immune evasion tool secreted by tumours to suppress cytotoxic immune cell activation and tumour destruction via an atypical, Wnt signalling independent pathway (Xiao et al., 2018). Nonetheless, it is a possibility that microglial-derived Dkk2 can also act upon microglia in an autocrine fashion at least in mice, although we can only speculate what the cellular response to such a stimulus would be.

Conversely, we provide evidence in support of a paracrine mechanism at least in cultured rat primary neurons as we demonstrate that recombinant human DKK2 protein blocks the synaptogenic effect of Wnt *in vitro*. Knowing that DKK2 can generally engage in Wnt antagonising and agonising activities depending respectively on the presence or absence of the co-receptor Kremen2 (Mao and Niehrs, 2003), it appears that, at least in our *in vitro* system, DKK2 protein acts as an antagonist. DKK2 may thus behave similarly to DKK1, a negative regulator of canonical Wnt/β-catenin and non-canonical Wnt/PCP signalling with known synapse destabilising properties (Purro et al., 2012; Galli et al., 2014; Killick et al., 2014; Marzo et al., 2016; Elliott et al., 2018; Sellers et al., 2018; see also review by Aghaizu et al., 2020), likely also in the human AD brain (Caricasole et al., 2004). Synapse density reductions in plaque proximal regions (Koffie et al., 2009) would be consistent with the fact that oligomeric βAmyloid induces *Dkk1* expression (Purro et al., 2012; Killick et al., 2014; Jackson et al., 2019). *Dkk2^+^*microglia accumulating around βAmyloid plaques may locally increase Dkk2 protein levels, adding to the anti-synaptic milieu established by Dkk1 near plaques. Given that microglia already engage in complement-mediated synaptic pruning by phagocytosis in AD mouse models (Hong et al., 2016; Shi et al., 2017), the relative contributions of individual synaptotoxic components around plaques will have to be addressed in future studies.

In assessing the chronological order between microgliosis/microglial plaque clustering and microglial *Dkk2* upregulation, we observed significant microgliosis increases before *Dkk2* upregulation in *APP/PS1*, *APP^NL-G-F^*, and *SOD1^G93A^* mice with respect to absolute quantification metrics (see Fig 1,2,4). However, when comparing relative rate changes, the rate of *Dkk2* signal increase at early disease stages in the *APP/PS1* AD mouse model surpassed the rate of microgliosis increase (Fig. 2*F,J*). It should be noted that *Dkk2* induction was initiated from near-zero basal expression levels (Fig. 2*E*), whereas both basal microglia numbers and Iba1 immunoreactivity levels were decidedly greater than zero (Fig. 2*C,D,G,H*). The potential for more pronounced changes was thus markedly greater for *Dkk2* induction at least in *APP/PS1* mice. Conversely, in S*OD1^G93A^* ALS mice, the rate of microgliosis increase surpassed that of *Dkk2* signal increase at early disease stages (Fig. 4*E*). Presumably, basal microglial cell densities lower than those observed in the mouse brain (Fig. 4*B,C* vs 2*C,D,G,H*), which is in keeping with published literature (Tan et al., 2020), likely contributed at least partially to this outcome.

What should be addressed in future studies is whether the ability to induce *Dkk2* expression is innate in all microglia or whether context, such as proximity to neurodegenerative lesions, is to be ascribed a more prominent role. CNS microglia are not a homogeneous population of cells, with gene expression signatures differing depending on factors such as brain region, sex, age, and context including disease (reviewed by Masuda et al., 2020). In 3 month old *APP^NL-G-F^* mice, *Dkk2^+^* ARM cells represented 6 % of to the total microglial pool (Sala Frigerio et al., 2019); this number increased to 33 and 52 % at 6 months and 12 months respectively. It will be interesting to discern whether ARM-competence is restricted to the initial population of ARM cells, which then serve as a proliferating seed population, or whether cells from the total microglia pool are continuously recruited into the *Dkk2^+^* ARM population as disease progresses.

Finally, our study has revealed discrepancies between human AD and transgenic AD mouse models. *DKK2* mRNA expression levels were not elevated in post-mortem frontal cortex samples from individuals diagnosed with AD vs healthy individuals. While other human brain and CNS regions like the spinal cord might exhibit *DKK2* upregulation (although unlikely given the absence of *DKK2* upregulation in recently published human RNA-seq databases; see below), the above finding is in stark contrast to our findings in neurodegeneration mouse models. A caveat worth mentioning is the fact that human microglia at AD end stage (Braak & Braak stage 5-6) were chronically exposed to disease for much longer periods than their mouse counterparts and chronic adaptations in microglia gene expression signatures may have masked potential earlier changes (we note that our pathological ageing samples at Braak & Braak stage 3-4 also lacked *DKK2* upregulation). Nevertheless, it is now known that gene expression signatures between mouse and human DAM/ARM populations, although overlapping to some extent, exhibit distinct differences (reviewed by Wang, 2021). In fact, numerous single-cell RNA-seq analyses have identified gene expression signatures that differed between mouse and human DAM/ARM populations (Grubman et al., 2019; Mathys et al., 2019; Nguyen et al., 2020; Olah et al., 2020; Smith et al., 2022). For technical reasons and in contrast to mouse studies, human single-cell RNA-seq studies are frequently, although not exclusively (Olah et al., 2020), restricted to nuclear transcripts, which may have contributed to the apparent transcriptomic differences between mouse and human microglia (note that extra- nuclear mRNA is abundant due to nuclear export before translation). However, even in a recent single nucleus RNA-seq comparative study involving human AD post-mortem tissue and the *5xFAD* AD mouse model, differences between human and mouse microglial gene expression signatures persisted (Zhou et al., 2020). The inability to detect extra-nuclear mRNA in human brain samples can be circumvented with the use of optimised tissue harvesting protocols (Olah et al., 2020), or *in situ* detection methods such as low throughput mRNA FISH (present study; Jolly et al., 2019) or higher throughput digital spatial profiling (Prokop et al., 2019). Nonetheless, our mRNA FISH based study strengthens the notion that human and mouse microglia, despite exhibiting some overlaps, are different even beyond just the expression status of *DKK2/Dkk2*, at least in the brain. Future studies should also examine any such interspecies differences in the spinal cord.

Our study therefore highlights the increasingly recognised difficulties and limitations of using mouse models to recapitulate facets of human biology and disease (Elder et al., 2010; Jucker, 2010; Cavanaugh et al., 2014; Justice and Dhillon, 2016; Perlman, 2016; Dawson et al., 2018). Regardless of whether this may be ascribed in our study to differing biological responses in humans vs mice or masking chronic adaptations in much longer human disease, these limitations likely play a key role in the absence of truly disease altering therapies to date despite decades of AD research and >100 clinical trials. Future AD research should thus substantially increase scrutiny in cases where animal models are to be used to ensure faithful modelling of human biology. Human based AD models including human induced pluripotent stem cell-derived cell cultures and brain organoids are potent additions to our tool-kit despite still lacking the capacity to fully recapitulate human *in vivo* biology in an *in vitro* setting, and indeed in an *in vivo* setting (Mancuso et al., 2019).

## Acknowledgements

We thank Dr. C.S. Frigerio for constructive discussions. This work was supported by the UK Dementia Research Institute, which receives its funding from DRI Ltd, funded by the UK Medical Research Council, Alzheimer’s Society and Alzheimer’s Research UK. LG and BK are supported by Brain Research UK and the Rosetrees Trust. We thank the Queen’s Square Brain Bank for providing human samples.

## Author contributions

(CRediT taxonomy) Conceptualisation, N.D.A., S.J., and P.J.W.; Methodology, N.D.A., S.J., S.K.S., B.K., and K.C.; Software, N.D.A. and S.J.; Validation, N.D.A. and S.J.; Formal Analysis, N.D.A.; Investigation, N.D.A. and S.J.; Resources, L.G., P.C.S., B.D.S., and P.J.W.; Writing – Original Draft, N.D.A.; Writing – Review & Editing, N.D.A., S.J., L.G., P.C.S., B.D.S., and P.J.W.; Funding Acquisition, L.G., P.C.S., B.D.S., and P.J.W.; Supervision, P.J.W.

---

**Supplementary Table S1.** mRNA FISH signal detection parameters. Related to Materials and methods, as well as Figure 6. Signal detection parameters used to identify *DKK2*, *TREM2*, and *P2RY12* mRNA FISH signal on confocal images form human samples using HALO software with the FISH-IF v2.0.4 module (Indica Labs).

|  | *DKK2* | *P2RY12* | *TREM2* |
| --- | --- | --- | --- |
| Contrast threshold | 0 | 0 | 0.71 |
| Signal minimum intensity | 0.13 | 0.13 | 0 |
| Spot size | 0.5, 20 | 0.5, 20 | 0.5, 20 |
| Copy intensity | 0.15 | 0.15 | 0.17 |
| Segmentation aggressiveness | 0.95 | 0.95 | 0.318 |

**Supplementary Table S2.** Human sample demographic data. Related to Figure 6. Table listing demographic data of individual subjects contributing to the generation of data set in Figure 6. Clinical presentation as well as post-mortem brain assessments are shown (brain weight, post-mortem (PM) delay, Braak & Braak stage, CERAD score, THAL stage, and ABC score).

| Sample ID (FCx) | Sample classification | Age at onset | Age at death | Duration (yrs) | Gender | Clinical presentation | Brain Weight (g) | PM delay (hrs:min) | Braak and Braak | CERAD | THAL | ABC score |
| --- | --- | --- | --- | --- | --- | --- | --- | --- | --- | --- | --- | --- |
| P48/15 | Control |  | 84 |  | M | Normal | 1468 | 79:10 | 0 | none | 0 | A0 B0 C0 |
| P58/10 | Control |  | 87 |  | F | Normal | 1114 | 51:40 | 1 | mild | 1 | A1 B1 C1 |
| P66/11 | Control |  | 86 |  | F | Normal/ path ageing | 1234 | 120:00 | 2 | none | 0 | A0 B1 C0 |
| P64/11 | Control |  | 80 |  | F | Normal/ path ageing | 1242 | 49:10 | 2 | none | 0 | A0 B1 C0 |
| P94/05 | Control |  | 71 |  | M | Normal/ Mild cerebrovascular disease | 1480 | 38:50 | 1 | none | 0 | A0 B1 C0 |
| P61/09 | Pathological aging |  | 99 |  | F | Normal/ path ageing | 1141 | 32:05 | 4 | mod dif/ mod mature | 3 | A2 B2 C2 |
| P5/05 | Pathological aging |  | 85 |  | F | Pathological ageing | 1263 | 39:10 | 3 | Mod dif/ Sparse mature | 3 | A2 B2 C1 |
| P12/10 | AD | 44 | 56 | 12 | F | Amnesic | 1034 | 53:00 | 6 | frequent | 5 | A3 B3 C3 |
| P73/08 | AD | 50 | 61 | 11 | M | Amnesic | 1372 | 54:10 | 6 | frequent | 5 | A3 B3 C3 |
| P88/10 | AD | 50 | 66 | 16 | F | Amnesic | 906 | 92:47 | 6 | frequent | 5 | A3 B3 C3 |
| P40/10 | AD | 51 | 62 | 11 | F | Amnesic | 978 | 62:55 | 6 | frequent | 5 | A3 B3 C3 |
| P71/10 | AD | 65 | 70 | 5 | F | Amnesic | 1233 | 46:58 | 5 | moderate | 5 | A3 B3 C3 |
| P76/11 | AD | 65 | 72 | 7 | M | Amnesic | 1325 | 38:55 | 5 | moderate | 5 | A3 B3 C3 |

